# A High-Throughput Assay to Identify Specific Nascent Chain Inhibitors

**DOI:** 10.64898/2026.09.03.749102

**Authors:** Patrick D. Fischer, Sebastian Hiller

## Abstract

“Undruggable” proteins without surface-accessible binding sites pose significant challenges to target-based drug discovery. Innovative approaches are needed to tackle these proteins. One promising strategy is targeting them in their nascent chain form at the ribosome, where they have a different conformation than in the folded form. A systematic approach to screen for such compounds has however been lacking. Here, we present a high-throughput assay to identify small molecules that specifically inhibit a protein of interest in its nascent-chain form. The assay employs a human *in vitro* transcription/translation system and monitors expression of the protein in real time via fluorescence detection. Specific inhibitors for the nascent chain of interest can then be identified by comparison with a counter screen. The assay was optimized to maximal sensitivity and reaction costs of ∼$0.01 per well, enabling large-scale screens. We validated performance with the reference compound PF846 on the nascent chain of the protein PCSK9. Feasibility for high-throughput screening was demonstrated using a library of 1,760 compounds against the oncogenic KRAS variant A146T and the protein ApoC3, with mean Z′ scores of 0.69 and 0.88, respectively. Sixteen global translation inhibitors were identified in each campaign, while no compounds met the criteria for POI-selective inhibition. The assay thus provides a robust platform for larger-scale screening campaigns.

## INTRODUCTION

Fundamental research in molecular biology and biomedicine including the completion of the human genome project has led to the identification of hundreds of drug targets, such as kinases, enzymes and receptors [1, 2]. A target is “druggable” if its activity can be modulated by small molecule drugs [3], and this comprises so far around 10% of the human proteome. Among these, around half are relevant for disease [4]. In contrast, a target that resists conventional approaches to identify inhibitors is – after some reasonable attempts – considered “undruggable” [5–7]. Quite often, such targets are found to feature flat and extended surfaces without well-defined binding pockets. Undruggable targets present fundamental challenges and require novel innovative approaches.

One such alternative approach is to target proteins in their nascent chain form, where they adopt a different structure than in their folded form. A small molecule that binds and inhibits the nascent chain in the ribosome exit tunnel leads to translation stalling and degradation of the protein before it ever reaches its active form. The principal feasibility of this approach has been demonstrated by the small molecule PF846, which was discovered in an inhibitor screen for proprotein convertase subtilisin/kexin type 9 (PCSK9) [8–12]. Detailed investigation revealed that the compound targets the nascent chain in a specific manner with few off-target effects. At first sight, it might seem counterintuitive that a nascent chain can be specifically inhibited. It is however possible, because the nascent chain in the ribosomal exit tunnel adopts unique conformations that can be targeted in a similar way as folded proteins. Whereas PF846 provided the proof-of-concept, no high-throughput screen for human ribosome nascent-chain complexes is currently available. A droplet DNA-encoded library (DEL) screen in wheat-germ extracts exists, demonstrating high-throughput screening potential [13], but this screen does not faithfully present the human ribosome and results are thus transferrable to human application in only very limited fashion.

The RAS gene family (KRAS, NRAS, and HRAS) plays a fundamental role in controlling cell proliferation and is consequently the most commonly mutated gene in many cancers [14]. The RAS proteins lack deep binding pockets, while featuring picomolar binding affinity to their natural ligands GDP and GTP, making them a prime example for an undruggable target. Recent efforts saw the development of allele-specific, covalent inhibitors of the KRAS mutant G12C, with promising clinical outcomes [15–19]. Alanine 146 is the fourth-most frequently mutated residue in KRAS, almost exclusively found in gastrointestinal or blood cancers [20–22]. KRAS A146T in particular is characterized by its increased nucleotide exchange rate and tissue-specific disruption of basal homeostasis [23]. In a cohort of patients with colorectal peritoneal metastases, A146T belonged to a KRAS mutation subgroup associated with a median overall survival of 31.2 months, compared with >120 months for a subgroup including G13C, Q61H and A146V [24]. Although progress is being made towards identifying specific non-covalent KRAS inhibitors [25] and pan-KRAS inhibitors [26], there is still an urgent need to identify small molecules targeting this oncogenic protein.

Here, we address the need for a robust high-throughput screen technology by developing a plate-format assay, enabling cost-effective screening against human nascent chains in human ribosomes. To this end, we use a coupled human in vitro transcription/translation (IVT/T) system with optimized fluorescent reporter and a matched counterscreen. Reaction composition and readout conditions were optimized to reduce reagent cost while maintaining robustness, allowing screening on standard plate readers in high-throughput format.

## RESULTS

### A fluorescent human in vitro transcription/translation system

We sought to develop an in vitro assay that reliably detects whether a specific mRNA is translated by human ribosomes. The assay should be sensitive to identify initial hits with weak affinity. We opted for a coupled in vitro transcription/translation (IVT/T) system, derived from human cell lysate, with fluorescence readout. Compared to an assay in cells, this approach is expected to minimize background interference. The system monitors synthesis of a fluorescent protein that is C-terminally fused to the protein of interest (Figure 1A). Small molecules that induce translational stalling prevent synthesis of the fluorescent tag and can therefore be identified by absence of fluorescence compared to a positive control (Figure 1B). We chose a coupled transcription/translation system to avoid the step of mRNA purification, thus enabling a simple, practical assay setup.

**Figure 1.**
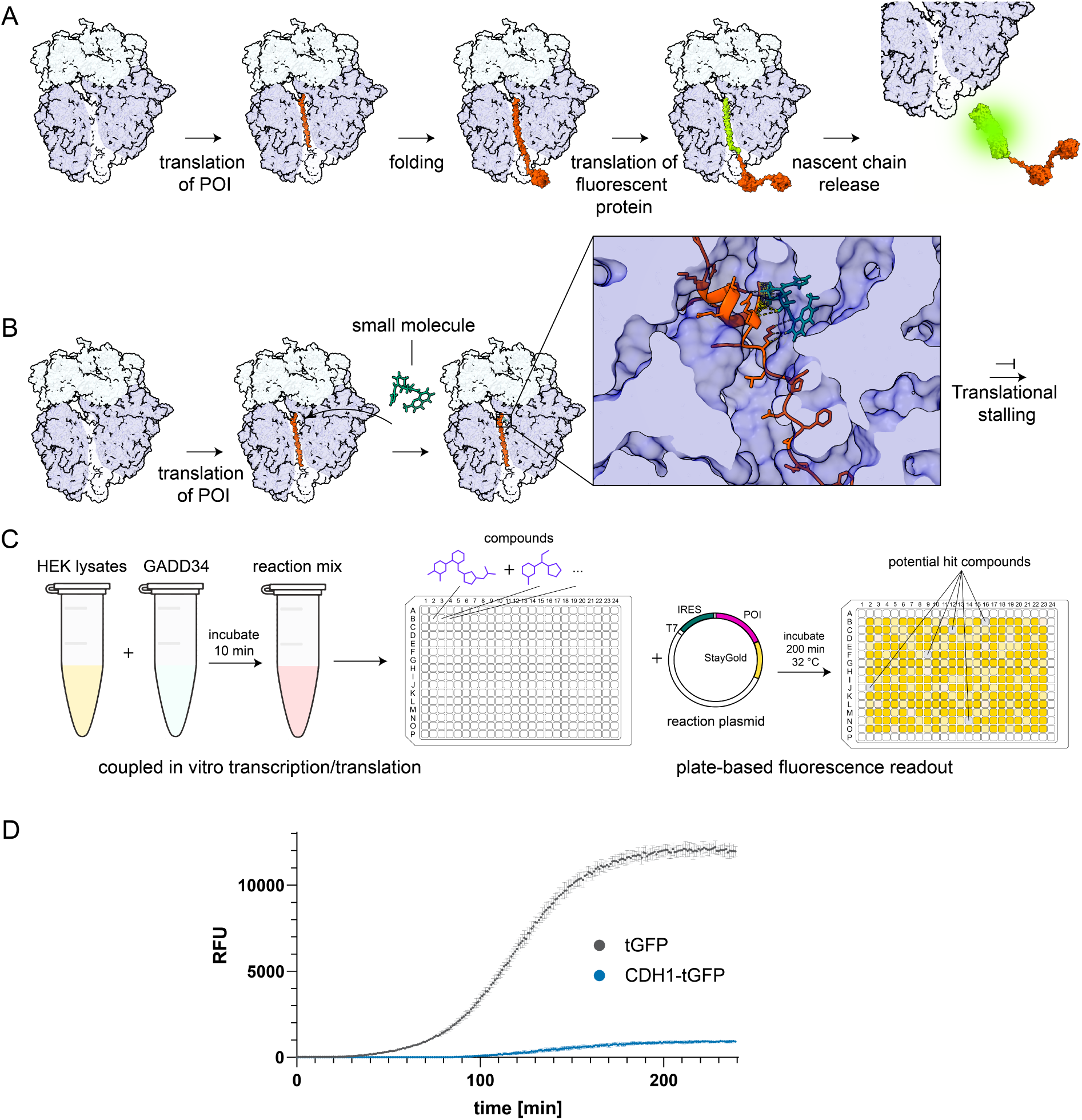
Assay concept. **A.** The protein of interest (POI) is translated with a C-terminal fluorescent tag. In absence of an inhibitor, a fluorescence signal emerges upon completion of translation. **B.** The presence of a translation inhibitor induces translational stalling and thus absence of the fluorescence signal. **C.** HEK cell lysates are incubated with the accessory protein GADD34 and then introduced to a reaction mix for coupled IVT/T reactions. The mixture is transferred to a 384 well-plate pre-plated with compound stocks and introduced to the reaction plasmid. Fluorescence signal is read out after 200 min of incubation at 32 °C. **D.** Reaction kinetics for the translation of tGFP and POI (CDH1)-tGFP.

The assay employs a DNA plasmid encoding the gene of interest under control of a T7 promoter. During the reaction, recombinant T7 RNA polymerase synthesizes the mRNA template, which contains an encephalomyocarditis virus (EMCV) internal ribosome entry site (IRES) to permit cap-independent translation (Figure 1C). The gene of interest is followed by a fluorescent reporter, and a T7 terminator defines the 3′ end. In the initial implementation of the assay, the fluorescent reporter was turboGFP (tGFP), a rapidly maturing fluorophore. In addition to the construct with the protein of interest (POI), the same compound library is counter screened against a fluorophore-only construct. This counterscreen removes global translation inhibitors and compounds that interfere with the fluorescent protein and its detection.

At the start of the reaction, the plasmid is incubated with human cell lysate, RNA polymerase, and a reaction mix (Table 1) in a typical volume of 10 µL. The reaction mix contains a set of seven accessory proteins that support transcription, translation, energy regeneration and maintenance of nucleotide pools [27–30]. This set includes the T7 RNA polymerase that is indispensable for mRNA synthesis, as well as accessory enzymes for energy regeneration and maintenance of nucleotide pools [31]. Recombinant factors GADD34 and K3L relieve translation inhibition caused by eIF2α phosphorylation [32–36], myokinase and creatine kinase regenerate ATP, nucleoside diphosphate kinase (NDK) regenerates GTP, and pyrophosphatase removes inhibitory pyrophosphate generated during transcription [31]. With this setup, fluorescence increased hyperbolically and reached a stable plateau, after 60–180 min. This kinetic profile is consistent with continuous synthesis and maturation of the fluorescent reporter with gradual loss of translational activity, confirming that signal accumulation reflects bona fide protein translation rather than nonspecific fluorescence fluctuations (Figure 1D).

**Table 1.** In vitro transcription/translation components and price optimization.

| Component | Price (USD/reaction) | % of total price | Optimized price (USD/reaction) |
| --- | --- | --- | --- |
| 20 amino acids | < 0.001 | < 0.01 | < 0.001 |
| HEPES-KOH | < 0.001 | 0.02 | < 0.001 |
| KOAc | < 0.001 | < 0.01 | < 0.001 |
| Mg(OAc)<sub>2</sub> | < 0.001 | < 0.01 | < 0.001 |
| glycerol | < 0.001 | < 0.01 | < 0.001 |
| spermidine | < 0.001 | 0.03 | < 0.001 |
| putrescine | < 0.001 | 0.13 | < 0.001 |
| DTT | < 0.001 | 0.01 | < 0.001 |
| ATP | < 0.001 | 0.26 | < 0.001 |
| GTP | 0.002 | 2.60 | 0.002 |
| CTP | 0.001 | 0.82 | 0.001 |
| UTP | 0.002 | 2.83 | 0.002 |
| creatine phosphate | 0.001 | 1.07 | 0.001 |
| creatine kinase | < 0.001 | 0.50 | < 0.001 |
| myokinase | 0.002 | 3.49 | 0 |
| NDK | 0.003 | 2.91 | 0 |
| pyrophosphatase | 0.001 | 2.16 | 0.001 |
| tRNA | (bovine) 0.050 | 71.92 | (*E. coli*) 0.004 |
| RNase inhibitor | 0.008 | 11.24 | 0 |
| T7 RNAP | recombinant | 0 | recombinant |
| GADD34 | recombinant | 0 | recombinant |
| K3L | recombinant | 0 | 0 |
| total | 0.069 | 100 | 0.011 |

The assay reached high sensitivity when expressing tGFP alone. However, when it was tested with cadherin 1 (CDH1_586-750_) as a protein of interest, it resulted in a fluorescence curve that increased and plateaued, with a sensitivity that was smaller than tGFP alone by a factor of 14 (Figure 1D). This difference likely reflects the additional biochemical demand of synthesis and increased probability of misfolding and aggregation with the longer fusion constructs, which reduces the effective reporter yield. The assay is thus functional, but the reduced sensitivity warranted further improvements.

### Optimization of sensitivity and cost

To make the assay suitable for high-throughput screening (HTS), we optimized the IVT/T reaction for both cost efficiency and sensitivity. We first optimized the reaction composition toward lower per-well costs (Table 1). Commercially available 1-step human coupled IVT/T kits exceed costs of $10/well and are therefore not suitable for high-throughput screening efforts [37]. Reconstitution of these formulations using the individual reagents according to standard literature reduces these costs to ∼$0.07/well [27–31, 38]. While this is a substantial economic improvement, we asked whether the reaction costs could be reduced further, as campaigns involving hundreds of plates and multiple assay conditions still result in significant reagent consumption.

As a first element for improvement, we recognized that tRNA supplementation is commonly required to maintain high translation activity, due to depletion of the endogenous tRNA pool during lysate preparation. Bovine tRNA is commonly used, because its sequence composition and modification pattern are expected to be compatible with the human translation machinery [31, 39]. However, bovine tRNA is also a dominant cost contributor. Strikingly, replacement of bovine tRNA with *E. coli* tRNA preserved full performance (Figure 2A) at ∼7 % of the cost, providing substantial savings. This finding is particularly remarkable, as the alternative substitution with yeast tRNA markedly decreased protein yield. These results indicate that efficient translation in the human lysate system depends on the source of exogenous tRNA, likely reflecting differences in tRNA composition and modification state. Next, we evaluated the need for each of the seven accessory proteins, as they represented major cost factors (T7 RNA polymerase, GADD34, K3L, myokinase, creatine kinase, pyrophosphatase, NDK). To assess each component’s necessity, we prepared reaction mixes omitting single elements and measured tGFP fluorescence. The results show that GADD34 is essential, whereas K3L and the energy-recovery enzymes myokinase and NDK did not significantly enhance translation (Figure 2B) and could hence be omitted. Also, the RNase inhibitor RNasin, which was another notable cost factor (∼11 % of total), could be omitted without loss of performance. Overall, our optimized formulation reduced the reagent cost for 5 mL of 5X reaction mix from $173.50 to $27.00, corresponding to approximately $0.01 per reaction well, thus rendering the assay economically suitable for large-scale screening.

**Figure 2.**
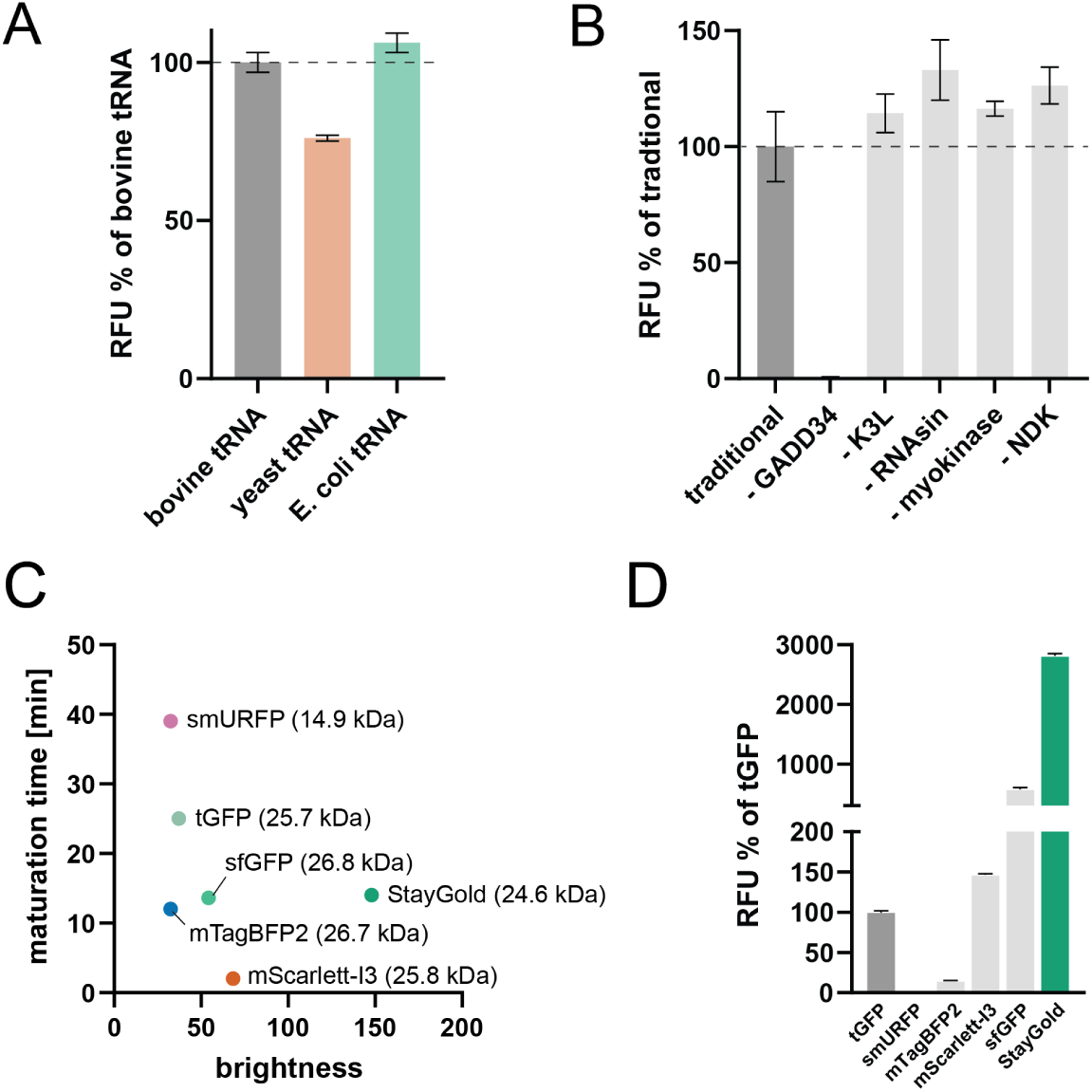
In vitro translation assay optimization. **A.** Performance of the human IVT/T assay with different exogenous tRNA, measured after 200 min using a tGFP-only reporter construct. Values are normalized to the bovine tRNA condition. **B.** Assessment of the effect of individual helper proteins on the performance of the human IVT/T assay, measured after 200 min using a tGFP-only reporter construct. Values are normalized to the complete reaction. Among the tested proteins, only GADD34 is indispensable. **C.** Properties of six fluorescent protein tags. **D.** Endpoint fluorescence of CDH1₅₈₆-₇₅₀ fused C-terminally to each fluorescent protein reporter, measured after 200 min under otherwise identical IVT/T conditions. Values are normalized to the CDH1₅₈₆–₇₅₀-tGFP construct.

We next aimed to optimize the type of the fluorescence reporter. We compared the properties of six different fluorescent proteins as C-terminal tags, using CDH1_586-750_ as the nascent chain. The six candidates were selected from the fluorescent protein database[40] based on their reported size, maturation time, lifetime, and brightness. These are key parameters influencing translation efficiency and detectable signal in vitro. The tested proteins were tGFP, smURFP, mTagBFP2, mScarlett-I3, sfGFP, and StayGold, representing a range of spectral classes and biophysical properties (Figure 2C). For each fusion construct, the fluorescence intensity was quantified after 200 min of IVT/T reaction (Fig. 2D). smURFP yielded no detectable signal, and mTagBFP2 reached only ∼15% of the tGFP signal. mScarlett-I3 moderately increased signal output (∼1.5-fold relative to tGFP), whereas sfGFP achieved a 5.7-fold improvement. The highest sensitivity was obtained with StayGold, which produced ∼28-fold higher fluorescence than tGFP. This sensitivity, together with its fast maturation time of 14 min, and exceptional photostability[41] make StayGold in our hands the best suited reporter for the IVT/T reaction. We therefore adopted StayGold as the fluorescence reporter in all subsequent assay implementations, providing a substantial increase in assay sensitivity while maintaining compatibility with the optimized low-cost reaction mix.

### Assay validation with the benchmark nascent chain inhibitor PF84C

Next, we sought to validate that the assay detects translation stalling and can quantify potency (IC_50_), by using the reported PCSK9 nascent chain inhibitor PF846. A fusion construct of PCSK9 including the stalling site was generated as PCSK9_1-148_-StayGold. Then, in vitro translation reactions of this construct were conducted in the absence and presence of PF846, titrated in the range of 6 nM to 1 mM. In our setup, PF846 inhibited PCSK9_1-148_-StayGold translation with an apparent IC_50_ of 27 µM (95% CI 7 - 40 µM) (Figure 3A). Previously reported IC_50_ values for PF846 vary between assay formats (∼0.3 µM in cell assays [10] and ∼4 µM in wheat-germ in vitro translation [13]). Importantly, our endpoint readout measures the amount of completed PCSK9-StayGold product accumulated over 200 min. A persistent translational stall therefore prevents reporter formation, whereas transient pausing followed by recovery can still result in reporter synthesis within the assay time. Likewise, slower translation can reduce reporter accumulation without complete arrest. These effects cannot be distinguished by the endpoint measurement and may, together with differences between translation systems, contribute to the higher apparent IC_50_ observed here. The benchmark response confirms that our coupled IVT/T assay sensitively reports sequence-dependent translation inhibition. Additionally, raw fluorescence distributions demonstrate that PF846 at maximal concentration reduces signal toward the no-template baseline, confirming robust assay window and clear separation between active inhibition and background signal (Fig. 3B).

**Figure 3.**
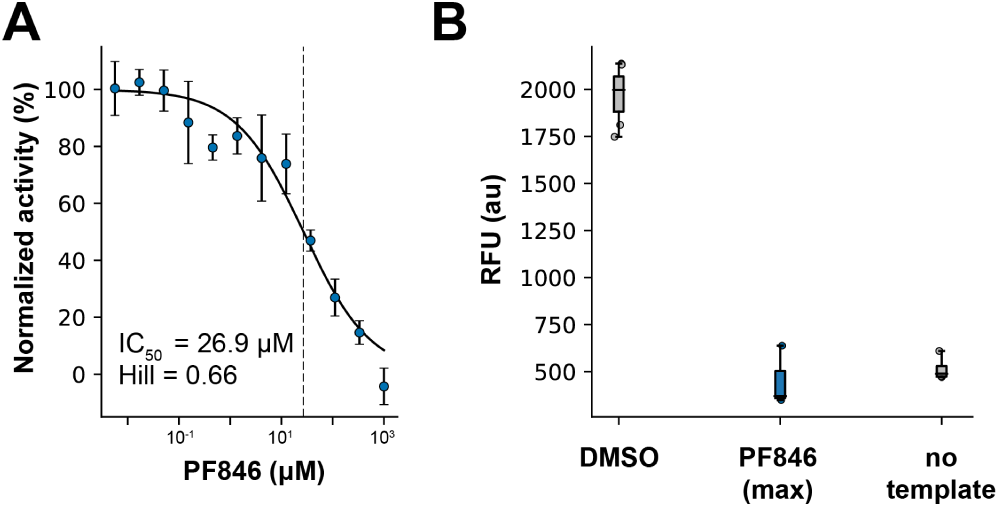
Proof of concept. **A.** IC50 determination of PF846 shows dose-dependent fluorescence reduction, with an apparent IC50 value of 27 µM. **B.** Comparison of absolute fluorescence endpoints for the DMSO controls, PF846 at the highest used concentration (1 mM) and the no-template control.

### Pilot screen against ApoC3 and KRAS A14CT

With a low-cost, high-sensitivity reporter and benchmark inhibitor response established, we next evaluated HTS performance in pilot screens against targets KRAS A146T and ApoC3. We screened a curated small molecule library of 1,760 compounds against the two proteins of interest, selected to include PF846-related chemistry and compounds with known or suspected effects on translation. The oncogenic KRAS mutant A146T was selected as a classical “undruggable” target to assess the applicability of the assay beyond established nascent-chain inhibitor systems. ApoC3 was selected as a distinct second target because of sequence similarity to PF846-sensitive nascent-chain sequences. The compound library included known pan-translation inhibitors, empty controls and PF846 analogues. Each construct was screened using five 384 well plates. Each plate contained 8 DMSO wells and no-template controls to define the assay floor. To account for evaporation effects, assay reactions for ApoC3 were performed using sealed plates, while KRAS A146T plates were left unsealed. Compounds were applied at a concentration of 50 µM. In addition to the two proteins of interest (POIs) with StayGold fusion, the counterscreen was performed in identical plates against StayGold only. Compound specificity was assessed by comparing POI-StayGold and StayGold-only responses. Formal hit classification was based on the Mahalanobis distance and directional Z score criteria described below.

As shown in Figure 4A, the ApoC3 campaign showed low variability in the positive DMSO controls, with CV values of 3.0-4.8% across plates and a mean CV of 3.9%. The DMSO CV describes the well-to-well variability of uninhibited translation reactions and is therefore an important measure of assay precision. No-template wells remained well separated from the DMSO distribution, resulting in HTS-grade Z′ values of 0.85-0.91, with a mean Z′ of 0.88. In addition to this Z′-based quality assessment, the assay showed a large dynamic range, with signal-to-background values of 215-359 and signal-to-noise values of 721-1586. These metrics describe how strongly the translation signal is separated from the no-template background and therefore indicate how robustly signal loss can be detected.

**Figure 4.**
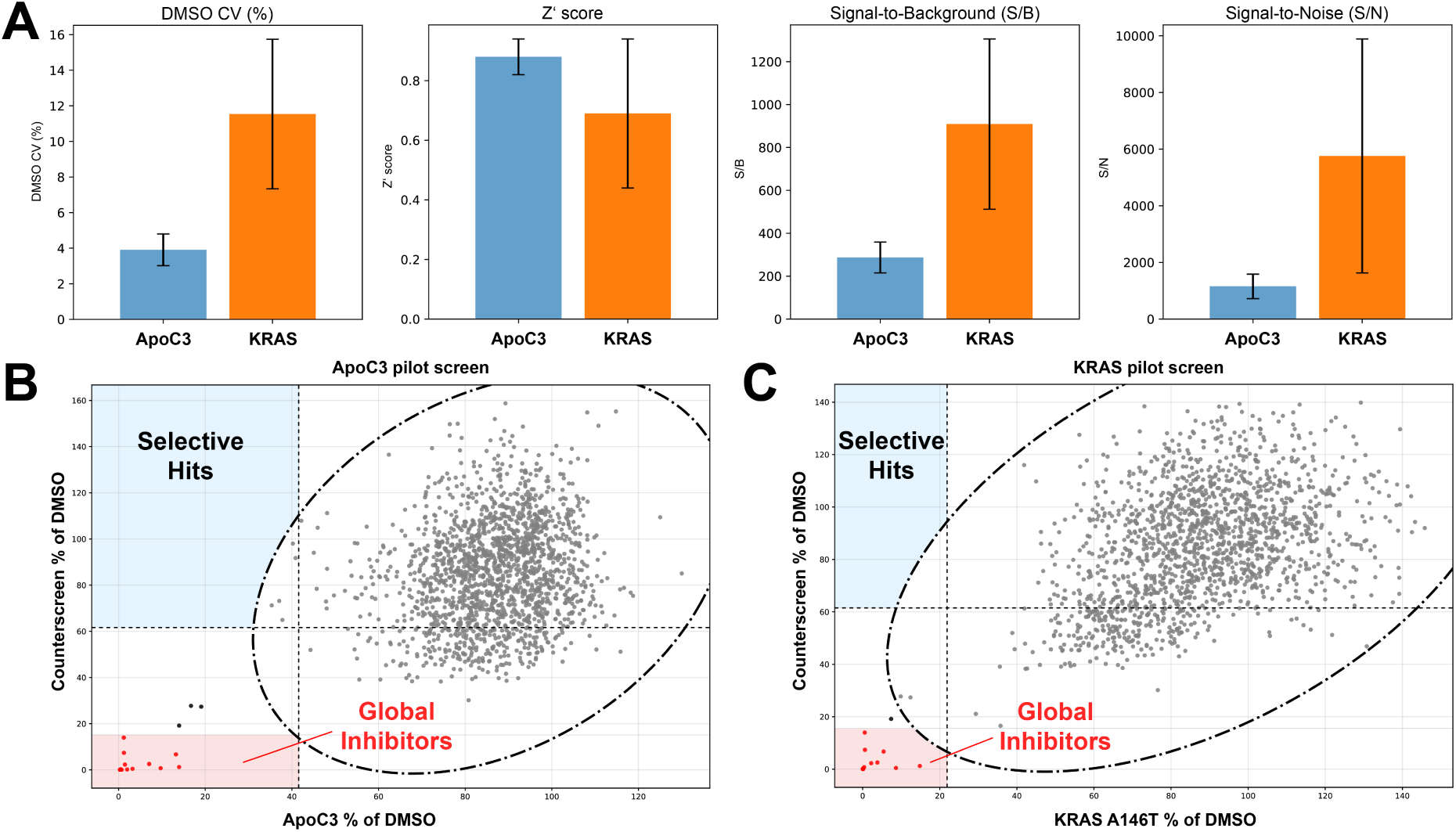
High-throughput quality control metrics and results of the two pilot screens. **A.** Comparison of DMSO variability, signal-to-background (S/B), signal-to-noise (S/N) and Z’ scores between ApoC3 and KRAS screening campaigns. **B-C.** Results of the ApoC3 and KRAS pilot screens. Compounds selectively inhibiting the POI appear in the top left corner and global translation inhibitors (red) in the bottom left corner.

In the KRAS A146T campaign, which was run on unsealed plates, DMSO variability was increased compared to the sealed ApoC3 campaign, with CV values of 7.3-16.0% and a mean CV of 10.0% (Figure 4A). This observation is consistent with increased sensitivity of the assay to evaporation. Despite this increased variance, KRAS A146T plates retained robust separation from no-template controls, with Z′ values of 0.53-0.78 and a mean Z′ of 0.69. The dynamic range also remained high, with signal-to-background values of 512-1306 and signal-to-noise values of 1627-9891. Z′ scores above 0.5 generally indicate robust assay performance for high-throughput screening [42], and control variability is generally considered acceptable at CV values below 20% and good below 10% [43, 44]. Thus, both campaigns met HTS-quality criteria, with the sealed ApoC3 campaign showing particularly high assay precision.

Next, we assessed compound behavior by plotting the POI and counterscreen responses against each other and then classifying compounds that were statistically distinct from the main compound population. To do so, we applied a Mahalanobis distance cutoff (MD ≥ 3.717) to identify compounds outside the 99.9% confidence ellipse of the compound cloud [45]. Compounds passing this outlier filter were then classified according to directional Z score cutoffs [46]: compounds with Z(POI) ≤ −3 and Z(counterscreen) ≤ −3 were classified as global inhibitors, whereas compounds with Z(POI) ≤ −3 and Z(counterscreen) > −1 were classified as putative POI-selective hits. Using this conservative framework, the ApoC3 campaign yielded 16 global inhibitors and no POI-selective candidates (Figure 4B), while the KRAS A146T campaign yielded the same 16 global inhibitors and no POI-selective candidates (Figure 4C). In addition, three ApoC3 compounds and one KRAS A146T compound showed a global-inhibitor-like response pattern but did not pass the Mahalanobis outlier cutoff. These compounds were therefore not included in the formal global-inhibitor count, but are best interpreted as borderline global inhibition rather than POI-selective activity.

## DISCUSSION

Targeting nascent polypeptides at the ribosome offers a conceptually attractive path to modulate proteins that remain difficult to inhibit in their folded states. However, discovery efforts have so far been constrained by the lack of broadly accessible screening formats that (i) operate in a human context, (ii) provide sufficient detection sensitivity, and (iii) are broadly applicable in multi-well plate format. Here, we establish a plate-based human IVT/T platform that addresses these bottlenecks through cost-optimization, a high-sensitivity StayGold endpoint readout, and a paired counterscreen design. Together, these elements support scalable screening on plate readers and thus routine implementation.

The assay was validated with the small molecule PF846, which is known to selectively stall human ribosomes on defined nascent-chain sequences [9, 11, 12]. Assay validation with PF846 provides an important benchmark because it is the best-characterized example of selective small-molecule engagement of the human ribosome exit tunnel leading to nascent-chain-dependent stalling. In our assay, PF846 produced a reproducible and concentration-dependent reduction in a PCSK9 construct, demonstrating that the fluorescence IVT/T workflow can detect a known nascent-chain-dependent stalling event. Furthermore, the ApoC3 and KRAS A146T pilot screens established assay robustness in 384-well format. Under the final hit-calling criteria, both campaigns identified the same number of global inhibitors (16 compounds each), indicating that the assay reproducibly detects compounds that suppress both POI and counterscreen output. In addition, a small number of compounds fell outside the main POI-counterscreen distribution without meeting the full Z score criteria for global inhibition. Importantly, no compounds fulfilled the final criteria for POI-selective inhibition in either screen. These results suggest that strongly target-selective translational stallers are rare, consistent with the expectation that sequence-selective stalling requires a specific combination of ribosome-binding chemistry and nascent-chain context.

A recent study described an activity-based DNA-encoded library (DEL) screen in which a wheat germ IVTT assay was miniaturized into picoliter microfluidic droplets and demonstrated compatibility with microfluidic droplet screening using a 5,348-member DEL [13]. In this work, compounds were released from library beads inside droplets, translation output was monitored with a GFP reporter, and selectivity was assessed with a PCSK9 frameshift counterscreen. That work highlights the power of microfluidics for massive throughput and coupling to DEL-based library formats. In contrast, the platform described here prioritizes accessibility and direct operation in a human translation environment using standard plate handling and readers, while embedding a paired counterscreen as a default analysis setup. Importantly, the two approaches are complementary: droplet microfluidics can unlock extraordinary scale, whereas a plate-based human IVT/T format lowers barriers to adoption and enables rapid iteration across constructs, counterscreens, and follow-up assays without requiring droplet instrumentation, bead-based compound display, or DEL decoding.

Several limitations and next steps follow directly from these observations. Fluorescence loss is an indirect proxy for translational arrest and can be influenced by fluorescence interference and upstream steps; therefore, any candidate requires orthogonal confirmation. In addition, the present screens used a single endpoint; incorporating simple kinetic features from the fluorescence time curves may help distinguish delayed expression from abrupt arrest and improve triage without sacrificing throughput. Rather than relying exclusively on endpoint measurements, simple features such as lag time, maximum signal accumulation rate, time to half-maximal signal, or total area under the curve could be extracted in a high-throughput manner. In principle, such parameters may help distinguish compounds that gradually delay reporter expression from those that cause a more abrupt arrest of translation, thereby adding an extra layer of triage without sacrificing the scalability of the assay. Finally, the absence of validated selective hits in this modest, curated library suggests that general screening collections may underrepresent chemotypes capable of engaging human ribosome nascent-chain complexes. Future efforts will likely require broader chemical diversification beyond the current PF846-focused space, together with systematic optimization for both ribosome engagement and nascent-chain-context selectivity.

## Methods

### Expression and purification of recombinant proteins

#### T7 RNA polymerase

His-tagged T7 RNA polymerase was expressed and purified according to [47], with the following alterations: Briefly, a plasmid coding for T7 RNAP in pT7-911Q was transformed into Lemo21(DE3) *E. coli* and plated on lysogeny broth (LB)-agar plates containing 100 μg/mL ampicillin. A preculture (10 mL) in LB containing 100 μg/mL ampicillin was inoculated with 1-5 colonies from this agar plate and grown at 37° C overnight. The next morning, 1 L of LB containing 100 μg/mL ampicillin was inoculated with this preculture and grown at 37° C until the OD_600_ reached 0.5-0.6. Protein expression was induced with 0.5 mM isopropyl β-D-1-thiogalactopyranoside (IPTG) and cells were grown for 3h at 37° C. After centrifugation at 6,000 x g for 15 min, the pellet was resuspended in 25 mL of lysis buffer (50 mM Tris-HCl pH = 8.0, 100 mM NaCl, 5% glycerol, 2 mM β-mercaptoethanol (BME), 10 mM imidazole) and 1.0 mL of 10 mg/mL lysozyme, 50 μL of 20 mg/mL PMSF, and 20 μL of 5 mg/mL leupeptin were added. After resuspension, 0.25 mL of 8% (w/v) sodium deoxycholate was added to the lysate and the suspension was incubated at 4° C for 1h. The cells were then lysed by sonication (35% amplitude, 1s on, 2s off) and centrifuged at 33,000 x g for 40 min at 4° C. The cleared lysate was incubated with ∼5 mL HisPure Ni-NTA resin (ThermoFisher) for 1h at 4° C. Then, the beads were washed with 10 column volumes (CV) lysis buffer. After washing, bound T7 RNAP was eluted with 10 CV elution buffer (lysis buffer with 500 mM imidazole). Eluted T7 RNAP was dialyzed against storage buffer (20 mM potassium phosphate pH = 7.5, 100 mM NaCl, 50% glycerol, 10 mM DTT, 0.1 mM EDTA, 0.2% NaN_3_), aliquoted and stored at -20° C.

#### GADD34

A construct coding for His6-TEV-GADD34240-674 in a pET28a vector was transformed into *E. coli* pLysS and plated on LB-agar plates containing 50 μg/mL kanamycin and 30 μg/mL chloramphenicol. A preculture (10 mL/L large culture) in LB containing 50 μg/mL kanamycin and 30 μg/mL chloramphenicol was inoculated with 1-5 colonies from this agar plate and grown at 37° C overnight. The next morning, 1 L of LB containing 50 μg/mL kanamycin and 30 μg/mL chloramphenicol was inoculated with this preculture and grown at 37° C until the OD_600_ reached 0.7. Protein expression was induced with 0.5 mM isopropyl β-D-1-thiogalactopyranoside (IPTG) and cells were grown for 3h at 37° C. After centrifugation at 6,000 x g for 15 min, the pellet was resuspended in ∼40 mL lysis buffer (50 mM Tris pH = 8.0, 350 mM NaCl, 20 mM imidazole, 2 mM BME, 10% glycerol, 5 mg lysozyme, 1 μL benzonase, 1 cOmplete™, Mini, EDTA-free Protease Inhibitor Cocktail [Roche]) per liter culture. The cells were then lysed by sonication (35% amplitude, 1s on, 2s off) and centrifuged at 33,000 x g for 40 min at 4° C. The cleared lysate was incubated with ∼5 mL HisPure Ni-NTA resin (ThermoFisher) for 1h at 4° C. Then, the beads were washed with 10 column volumes (CV) lysis buffer (without lysozyme, benzonase and protease inhibitors). After washing, bound GADD34 was eluted with 10 CV elution buffer (lysis buffer with 350 mM imidazole). Eluted GADD34 was dialysed after the addition of TEV protease against 50 mM HEPES pH 7.4, 150 mM KOAc, 5 mM Mg(OAc)_2_, 10% glycerol, 2 mM DTT overnight at 4° C. The protein was further purified using size exclusion chromatography (HiLoad Superdex 200 pg, Cytiva) preequilibrated with dialysis buffer, aliquoted and stored at -80° C.

#### K3L

A construct coding for His6-TEV-K3L in a pET28a vector was transformed into E. coli pLysS and plated on LB-agar plates containing 50 μg/mL kanamycin and 30 μg/mL chloramphenicol. A preculture (10 mL/L large culture) in LB containing 50 μg/mL kanamycin and 30 μg/mL chloramphenicol was inoculated with 1-5 colonies from this agar plate and grown at 37° C overnight. The next morning, 1 L of LB containing 50 μg/mL kanamycin and 30 μg/mL chloramphenicol was inoculated with this preculture and grown at 37° C until the OD_600_ reached 0.4. The temperature was reduced to 20° C and the culture was grown until the OD_600_ reached 0.7, when protein expression was induced with 0.5 mM isopropyl β-D-1-thiogalactopyranoside (IPTG). Cells were grown at 20° C overnight and harvested at 6,000 x g for 15 min the next morning. The pellet was resuspended in ∼40 mL lysis buffer (20 mM HEPES pH = 7.5, 100 mM KCl, 20 mM imidazole, 2 mM BME, 10% glycerol, 0.1 % Triton-X, 5 mg lysozyme, 1 μL benzonase, 1 cOmplete™, Mini, EDTA-free Protease Inhibitor Cocktail [Roche]) per liter culture. The cells were then lysed by sonication (35% amplitude, 1s on, 2s off) and centrifuged at 33,000 x g for 40 min at 4° C. The cleared lysate was incubated with ∼5 mL HisPure Ni-NTA resin (ThermoFisher) for 1h at 4° C. Then, the beads were washed with 10 column volumes (CV) lysis buffer (without lysozyme, benzonase and protease inhibitors). After washing, bound K3L was eluted with 10 CV elution buffer (20 mM HEPES, pH = 7.5, 150 mM KCl, 10% glycerol, 2 mM BME and 250 mM imidazole). Eluted protein was loaded onto a Mono Q Anion exchange column (Cytiva) preequilibrated with Buffer A (20 mM HEPES, pH = 7.5, 100 mM KCl, 10% glycerol, 0.1 mM EDTA and 5 mM BME). Bound protein was eluted with a gradient of Buffer B (20 mM HEPES, pH = 7.5, 500 mM KCl, 10% glycerol, 0.1 mM EDTA and 5 mM BME) and further purified by size exclusion chromatography (HiLoad Superdex 75 pg, Cytiva) preequilibrated with storage buffer (50 mM HEPES pH 7.4, 150 mM KOAc, 5 mM Mg(OAc)_2_, 10% glycerol, 2 mM DTT), aliquoted and stored at -80° C.

#### Cloning

All cloning procedures were performed using PCR conditions from IVA cloning and primers were designed using https://ivaprime.com. [48, 49] All constructs were cloned in a pUC-type vector backbone with Ampicillin resistance and the following elements (in order): A T7 promoter sequence, a 49 bp linker, an EMCV-IRES element, an ATG start codon, 3xFLAG (DYKDDDDK), a sequence coding for the POI, a sequence coding for turboGFP (or other indicated fluorescent proteins), a TGA stop codon, a 30xA poly-A-tail and a T7 terminator sequence.

### In vitro transcription/translation

#### Cell lysate preparation

HEK 293 suspension cells were cultured at 37° C in 293 SFM II medium (+ GlutaMAX). Cells were harvested at 0.8-1.0 × 10^6^ cells/mL at 500 x g for 5 min at room temperature. The pellet was washed 3 times with washing buffer (35 mM HEPES-KOH pH 7.5, 140 mM NaCl, 11 mM glucose) using 6 ml buffer per wash per mL pellet. After, cells were washed once with extraction buffer (20 mM HEPES-KOH pH 7.5, 45 mM KOAc, 45 mM KCl, 1.8 mM Mg(OAc)_2_, 1 mM DTT). Then, the pellet was resuspended in extraction buffer (to a final concentration of 3 x 10^8^ cells/ml, ∼ 1:1 pellet to buffer). Cells were lysed by shearing through a 26G needle ∼ 10 times. After this, the lysate was mixed with 1/29th high potassium buffer (20 mM HEPES-KOH pH 7.6, 945 mM KOAc, 945 mM KCl, 1.8 mM Mg(OAc)_2_, 1 mM DTT). The lysate was centrifuged twice at 1,200 x g for 5 min at 4° C. Aliquoted lysates are stored at a concentration of 25-30 mg/ml total protein concentration.

#### Reaction mix composition

The initial reaction mix was prepared as a 5X stock solution using the following ingredients:

260 mM HEPES-KOH, pH 7.4, 175 mM KOAc, 20 mM Mg(OAc)_2_, 20 μl/ml amino acid mix (1 part 50 X essential amino acids solution (Gibco), 1 part 100 X non-essential amino acids solution (Gibco), 1 part 20 mM L-glutamine), 5% glycerol, 2.75 mM spermidine, 3.5 mM putrescine, 25 mM DTT, 6.25 mM ATP, 4.15 mM GTP, 4.15 mM CTP, 4.15 mM UTP, 100 mM creatine phosphate, 300 ug/ml creatine kinase, 23.25 μg/ml myokinase, 2.4 μg/ml nucleoside-diphosphate kinase, 2 units/ml pyrophosphatase, 0.5 mg/ml Calf liver tRNA, 10 μl/ml RNasin, 550 nM T7 RNA polymerase.

The optimized reaction mix was prepared as a 5X stock solution using the following ingredients:

260 mM HEPES-KOH, pH 7.4, 175 mM KOAc, 20 mM Mg(OAc)2, 20 μL/mL amino acid mix (1 part 50X essential amino acids solution (Gibco), 1 part 100X non-essential amino acids solution (Gibco), 1 part 20 mM L-glutamine), 5% glycerol, 2.75 mM spermidine, 3.5 mM putrescine, 25 mM DTT, 6.25 mM ATP, 4.15 mM GTP, 4.15 mM CTP, 4.15 mM UTP, 100 mM creatine phosphate, 300 μg/mL creatine kinase, 2 units/mL pyrophosphatase, 0.5 mg/mL *E. coli* tRNA and 550 nM T7 RNA polymerase.

The ingredients together with RNase-free water are mixed and the reaction mix is aliquoted and stored at -80° C.

#### Coupled in vitro transcription/translation reaction

Coupled in vitro transcription/translation reactions are started in the following way:

5 μL lysate is incubated for 10 min at room temperature with 0.5 μL of GADD34 (10 μM) and/or 0.5 μL of K3L (10 μM) in a 384-well plate (Corning 3820: 384 Well Low Volume Fluorescence Plate). Then, 2 μL 5X reaction mix, 1-1.5 μL RNase-free water and 1 μL of plasmid DNA (100 ng/μL) are added and the reaction is initiated at 32° C.

##### Fluorescence readout

The fluorescence readout of the coupled in vitro transcription/translation reaction is performed using a Tecan Spark Multi-Mode Plate-Reader. The temperature was set to 32 °C, and plates were shaken in double-orbital mode for 5 s at an amplitude of 1 mm and 270 rpm. Monochromator settings were as follows:

*GFP:* Excitation 488 nm, bandwidth 5 nm, emission 510 nm, bandwidth 5 nm.

*smURFP:* Excitation 642 nm, bandwidth 5 nm, emission 670 nm, bandwidth 5 nm.

*mTagBFP2:* Excitation 399 nm, bandwidth 5 nm, emission 454 nm, bandwidth 5 nm.

*mScarlett-I3:* Excitation 568 nm, bandwidth 5 nm, emission 592 nm, bandwidth 5 nm.

*StayGold:* Excitation 493 nm, bandwidth 5 nm, emission 510 nm, bandwidth 5 nm.

Signal was recorded with a gain of 150, using 30 flashes and an integration time of 40 μs every minute.

### High-throughput assays

#### Plate layout, controls and compound screening

All screening experiments were performed in black polystyrene, flat bottom 384-well plates (Corning). Each assay plate contained compound wells, 8 DMSO control wells to define the plate reference signal and 8 no-template control wells to define the assay floor. No-template controls were prepared identically, except that the plasmid DNA template was omitted and replaced by an equal volume of nuclease-free water. For all analyses, DMSO and no-template wells were treated as plate-specific controls and were excluded from compound hit calling. Counterscreen plates were prepared and handled in the same 384-well format and used the same control layout. The counterscreen construct encodes StayGold alone (no POI sequence) and was assayed in parallel under otherwise identical conditions.

Compounds were screened at a single-point concentration of 50 µM in a total reaction volume of 10 µL per well in 384-well plates. Compounds were transferred from stock plates using an Echo Acoustic Liquid Handler (Beckman Coulter), and the final solvent concentration was kept constant across all compound and control wells. DMSO control wells contained the same final concentration of DMSO as compound wells (final DMSO: 0.5% v/v). Lysates, reaction mix, compounds and DNA templates were mixed by pipetting up and down carefully (Integra Evolve 16-channel pipettes). For the paired counterscreen workflow, each compound plate was run in parallel against (i) the POI–StayGold fusion construct and (ii) the StayGold-only counterscreen construct. Plates were handled identically, including control layout, reaction volumes, incubation conditions, and readout settings. For the ApoC3 campaign, plates were sealed using AlumaSeal® 384 film (Sigma Aldrich) to minimize evaporation; KRAS A146T plates were left unsealed as indicated in the main text.

Compound pre-incubation was performed with lysate for 15 min at room temperature prior to initiating the IVT/T reaction by addition of DNA template and reaction mix.

#### Data processing and quality-control metrics

Raw fluorescence values were read at the endpoint timepoint (200 min). For screens and follow-up assays, fluorescence values were normalized plate-wise to the mean of the DMSO controls of the corresponding plate to obtain “% of DMSO” values. For each plate, assay precision and separation were quantified using standard HTS metrics:

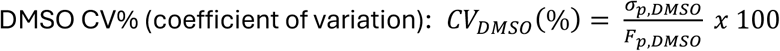

with σp,DMSO = standard deviation of the DMSO wells on plate p and Fp,DMSO = raw fluorescence of the DMSO wells on plate p.

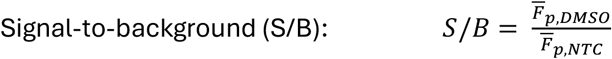

with *F̅_p,NTC_* = mean of the no-template control wells

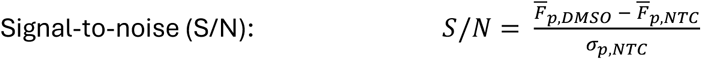

With σp,NTC = standard deviation of the no-template wells.

Z′ factor:

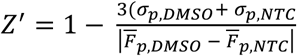

Z′ was computed plate-wise using DMSO wells as the “positive” control distribution and no-template wells as the “negative” control distribution. Unless otherwise stated, QC metrics were computed for the POI channel; counterscreen QC values were computed analogously. No outlier removal was applied to DMSO or no-template wells unless explicitly stated.

#### PF84C dose-response analysis and IC₅₀ calculation

Dose–response experiments with PF846 were performed in triplicate at the indicated concentrations. Compounds were preincubated with lysate for 15 min prior to initiation of the IVT/T reaction. Endpoint fluorescence values were normalized to the mean of DMSO controls on the same plate to obtain normalized activity (% of DMSO). Normalized dose–response data were fitted using a four-parameter logistic (4PL) model of the form:

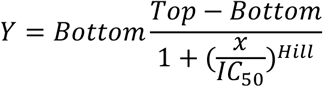

Where Y is the normalized response (% of DMSO), x is the compound concentration, Top and Bottom represent the upper and lower plateaus, IC50 is the apparent half-maximal inhibitory concentration and Hill is the Hill slope. Curve fitting was performed using nonlinear least squares regression in Python (SciPy curve_fit). Unless otherwise specified, the top plateau was constrained to 100% of DMSO. Where appropriate, the bottom plateau was constrained based on the mean no-template signal to reflect the assay floor. Apparent IC₅₀ values are reported with 95% confidence intervals derived from the regression fit.

## Acknowledgements

This work was funded by the EMBO Postdoctoral Fellowship (ALTF 707-2022) to P.D.F. and the Swiss National Science Foundation (CRSII5_209252). We thank Prof. Dr. Haribabu Arthanari for supplying a compound library for initial testing and Novartis for providing the focused compound library used for screening.

## Author contributions

P.D.F. and S.H. conceived and designed this work. P.D.F. performed the experiments and analysis. P.D.F. and S.H. wrote and revised the manuscript.

## Data availability

The data underlying this article are available from the corresponding authors upon reasonable request.

